# HIV-1 Vpr-induced Proinflammatory Response and Apoptosis are Mediated through the Sur1-Trpm4 Channel in Astrocytes

**DOI:** 10.1101/2020.03.19.999268

**Authors:** Ge Li, Tapas Makar, Volodymyr Gerzanich, Sudhakar Kalakonda, Svetlana Ivanova, Edna F. R. Pereira, J. Marc Simard, Richard Y. Zhao

## Abstract

There are about 38 million people currently living with HIV/AIDS worldwide. Successful treatment with combinational antiretroviral therapies (cART) can eliminate active replicating viruses and prolong lives to nearly normal lifespans. However, the new challenge faced by more than half of those HIV-infected and aging patients is chronic CNS neuroinflammation, which leads to HIV-associated neurocognitive disorders (HAND). While severe and progressive HAND has decreased significantly due to cART, chronic HAND often persists, resulting in high rates of delirium, dementia and depression that could lead to suicide. Indeed, the risk of suicide mortality in HIV-infected persons is significantly higher than in HIV-uninfected counterparts. Nevertheless, the mechanism of neuropathogenesis underlying HAND is not well understood. HAND is typically characterized by HIV-mediated glial neuroinflammation and neurotoxicity. Interestingly, the severity of some HAND does not always correlate with the levels of HIV, but rather with glial activation, suggesting other HIV-associated factors, not the whole virus *per se*, contribute to those HAND. HIV-1 viral protein R (Vpr) might be one of those viral factors, because Vpr induces neuroinflammation and causes neuronal apoptosis. The **objective** of this study was to delineate the specific role(s) of Vpr in activation of host neuroinflammation and neurotoxicity, as well as its contribution to HAND.

In this report, we show correlations between HIV expression and activation of proinflammatory markers (TLR4, TNFα, and NFκB) and the Sur1-Trpm4 channel in astrocytes of HIV-infected postmortem human and transgenic mouse brain tissues. We further show that Vpr alone activate the same set of proinflammatory markers in an astrocytic cell line SNB19. Vpr-induced host cell proinflammatory responses result in apoptotic cell death. Together, our data suggest that HIV-1 Vpr-induced proinflammatory response and apoptotic cell death are mediated through the Sur1-Trpm4 channel in astrocytes.

## Introduction

There are about 38 million people currently living with HIV/AIDS worldwide. Successful treatment with combinational antiretroviral therapies (cART) can eliminate active replicating viruses and prolong patients’ lives to nearly normal lifespans. However, the new challenge faced by many aging patients is chronic neuroinflammation in the central nerve system (CNS), leading to various HIV-associated neurocognitive disorders (HAND). More than 50% of HIV-infected individuals are afflicted with HAND. While severe and progressive HAND has decreased significantly due to cART, mild-to-moderate neurocognitive impairment often persists, resulting in high rates of delirium, dementia and depression that can lead to suicide^2^. Indeed, the risk of suicide mortality in HIV-infected persons is significantly higher than in HIV-uninfected counterparts^3,4^. xNevertheless, the mechanism of neuropathogenesis underlying HAND is not well understood. Glial cells such as microglia or astrocytes are thought to be the primary host innate immune effectors in HIV CNS infection, which triggers proinflammatory responses and activates neurotoxic astrocytes to kill neurons^5^. Thus, glia play a vital role in chronic neuroinflammation and neurotoxicity that is responsible for various manifestations of HAND, which includes not only progressive neurocognitive disorders, but also residual neurological impairment^6^. Chronic HAND is characterized by glial activation, cytokine/chemokine dysregulation, and neuronal damage and loss. Interestingly, the severity of some HAND types is not directly correlated with the level of HIV replication or viral load, but rather with glial activation^7^, suggesting that other HIV-associated factors, not the whole virus *per se*, contribute to HAND.

HIV-1 viral protein R (Vpr) might be one of those HIV-1 proteins contributing to HAND, because: 1) in the absence of active HIV-1 viral replication under cART, Vpr can still be detected in the circulation as a soluble and free protein^8^; 2) it is a transducing protein that can be taken up by glia and neurons^9^; 3) it is a neurotoxin that induces neuronal apoptosis^10^; 4) it binds to HIV-1 LTR promoter triggering viral transcription in latently-infected cells^11^; and 5) its effects are linked to various types of HAND^12,13^. In spite of these evidence indicating a prominent role of Vpr in HAND, how exactly Vpr contributes to HAND remains elusive. The objective of this study was to study the specific role(s) of Vpr in host innate proinflammatory responses, neurotoxicity and its contribution to HAND.

## Materials and Methods

### Cell culture and tissues

SNB19 (ATCC-CRL-2219), provided by Dr. HL Tang ^14,15^, is a human glioblastoma cell line, which was maintained in RPMI 1640 medium supplemented with 10% fetal bovine serum (FBS, Invitrogen) and 100 units/mL penicillin plus 100 µg/mL streptomycin. Postmortem human brain tissues from HIV-infected and non-infected individuals were from the University of Maryland Baltimore Brain and Tissue Bank. HIV-transgenic (Tg26) FVB/N mouse contains a partial pNL4-3 HIV-1 genome of 7.4 kb that encompasses most of the gag and *pol* genes including the *vpr* gene that been described previously (Kopp, 1992). The hippocampal and cortex regions of the brain sections were used for the immunohistochemistry (IHC) straining.

### Adenoviral constructs and cell transduction

The Adenoviral (Adv) Vpr construct was generated in this laboratory ^16,17^ by using the AdEasy Adenoviral Vector system (Cat#: 240009; Stratagene, La Jolla, CA). The viral titer of the Adv control was 2 × 10^9^ plaque forming unite (pfu)/mL, and the Adv-Vpr was 1 × 10^9^ pfu/mL, which were determined by an ELISA Adeno-X Rapid Titer Kit (Cat#: 631028, Takara). It detects the Adenoviral Hexon surface antigen. For Adv transduction, SNB19 cells in the concentration of 1 × 10^4^/well in 96 well plates were seeded and incubated at 37 C°/5% CO2 overnight to allow the cells to attach to the wells. The second day, SNB19 cells were transduced with Adv with the MOI as indicated. The Adv transduced cells were incubated at 37°C with gentle agitation. The cells were collected at indicated times for analyses.

### Immunostaining of mouse brain tissues

Paraffin-embedded Tg26 mouse brain sections (7 μm) were deparaffinized and rehydrated by passing through xylene and alcohols (100%, 95%, 70%). After antigen unmasking step, samples were incubated in blocking solution (90% TBST and 10% FBS) for 1 h, in primary antibody solutions for overnight, in secondary antibody solutions for 45 mins, in avidin-biotin complex (ABC) for 30 mins and finally, in 3,3’-diaminobenzidyne (DAB) solution until the expected color change is seen. After the staining step, slides were examined by using a light microscope at 40x. Two, three, or four slides were used for cell count in each group. Images were taken from both the cortex and the hippocampus areas. Five images were taken from each cortex, whereas whole cells of hippocampus were analyzed. The cells stained with the DAB substrate were taken as positive cells. Statistical Bonferroni’s Multiple Comparison Post Test was used for one-way ANOVA using Prism software (GraphPad, San Diego, CA). Statistical significance was accepted at the 95% confidence level (p < 0.05).

### Real-time RT-PCR

Total RNA was extracted from SNB19 cell line or snap-frozen mouse brain tissue by using TRIzol reagent (Life technologies) according to the manufacturer’s protocol. For snap-frozen mouse brain tissue, tissues were homogenized by Precellys Evolution Tissue Homogenizer (Bertin Technologies) then total RNA was extracted by using TRIzol reagent. RNA pellet was resuspended in RNase-free distilled water and stored at -80°C. 500 ng of RNA was used for Realtime RT-PCR analysis using iTaq universal SYBR Green One-Step Kit (BioRad) according to the manufacturer’s instructions. The primer sequences targeting interested genes were listed in attached table. Gene amplification in BioRad CFX96 real-time PCR system involved reverse transcription reaction at 50°C for 10 min, activation and DNA denaturation at 95°C for 1 min followed by 40 amplification cycles of 95°C for 15 s and 60°C for 30 s. mRNA expression (fold induction) was quantified by calculating the 2^-ΔCT^ value, with glyceraldehyde-3-phosphate dehydrogenase (GAPDH) mRNA as an endogenous control.

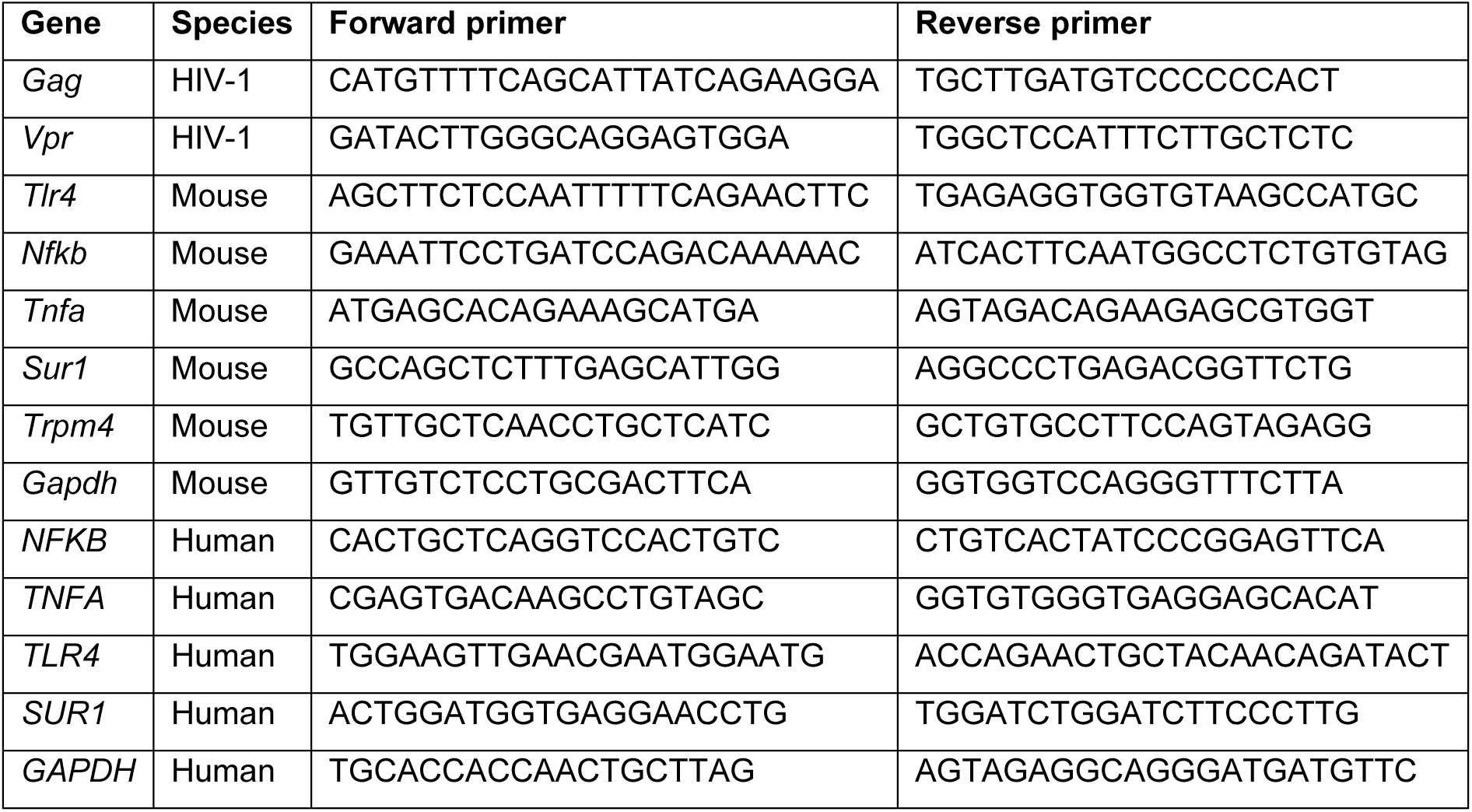

### Measurement of HIV-1 Vpr-induced Cellular Necrosis and Apoptosis

Cellular necrosis and apoptosis were measured by a RealTime-Glo™ Annexin V Apoptosis and Necrosis Assay kit (Promega) according to instruction of manufacturer. Briefly, 1 × 10^4^ cells/well were seeded into a 96-well plate and cultured at 37°C/5% CO2 overnight. After viral transduction with adenovirus, add 2x detection reagent (which included Annexin V NanoBiT® substrate, CaCl2, Necrosis Detection Reagent, Annexin V-SmBiT and Annexin V-LgBiT) into each tested well of 96-well plate. Incubate at 37°C/5% incubator and record luminescence (RLU) and fluorescence (RFU, 485nmEx/520–30nmEm) measurements at the desired time points with a SYNERGY-H1 microplate reader.

## Results

### HIV-1 infection elicits proinflammatory responses (TLR4, TNFα and NFκB) and upregulates Sur1 and Trpm4 in astrocytes of Tg26 mouse brain tissues

The **goal** of this experiment was to evaluate the impact of HIV-1 infection on neuroinflammation in the brain of HIV-infected mice. A transgenic (Tg) mouse line (Tg26)^18^ was used because of its clinical relevance to cART and continuous stress posed by HIV and viral proteins^19^. Brains of Tg26 and control WT mice (both males and females, 30±6 mg, 16 weeks old) were fixed with formalin and paraffin embedded (FFPE) for immunohistochemistry staining (IHC) using VECTASTAIN ABC Kits (Vector Laboratories, Burlingame, CA). As shown in **Fig. 1**, three proinflammatory regulators TLR4, NFkB and TNFα were significantly elevated (*, *p*<0.05; ***, p<0.001) in HIV-infected hippocampus in comparison with the WT controls. The levels of TLR4, NF-kB and TNFα elevated in parallel with the upregulation of glial fibrillary acidic protein (GFAP), a marker for astrocytes. Interestingly, both Sur1 and Trpm4 proteins were also highly elevated (***. *p*<0.001), which together form Sur1-Trpm4 channels that regulate proinflammatory responses^20,21^.

**Fig. 1.**
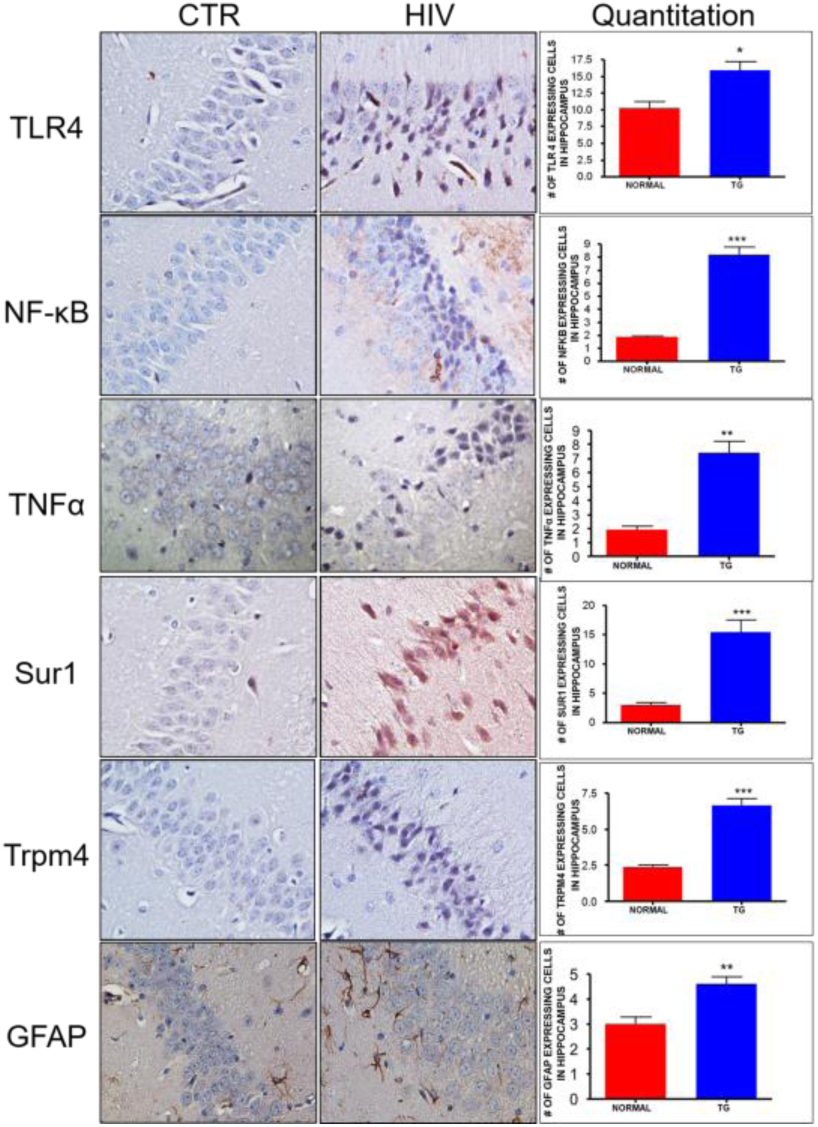
Correlation of HIV-1 infection with host innate immune proinflammatory responses in the hippocampus of the Tg26 mice. Each panel shows immunohistochemical straining of hippocampal sections with respective protein expressing cells in the wildtype control (CTR, left), Tg26 mice (HIV, middle). Graph: data (X±SD) were obtained from 3 mice/group. All images are 40x in magnification. Two-tailed and paired t-test was used for statistical comparison analysis: *, *p*<0.05; **, p<0.01; ***, p<0.001.

To verify the observed findings, RT-qPCR was used for mRNA transcriptional analyses of the same mouse brain tissues. The HIV-1 infection status of Tg26 mice was confirmed by the presence of *gag* and the *vpr* gene transcripts in comparison with the WT controls (**Fig. 2**, insert). Additional testing showed a positive correlation between gene transcription of HIV-1 *gag*/*vpr* and *TLR4, TNFα* and *NFκB*, as well as with *Abcc8*/Sur1 (**Fig. 2**).

**Fig. 2.**
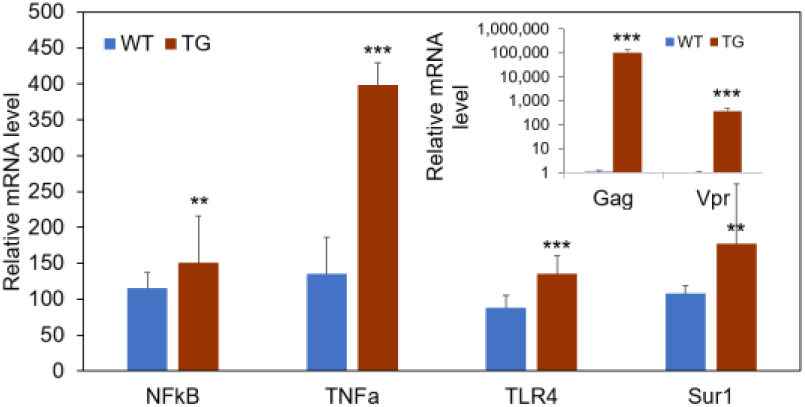
mRNA transcriptional analysis by RT-Qpcr shows correlation of HIV and Vpr with host innate proinflammatory responses in the brain tissues of Tg26 mice. Total mRNA was extracted from mouse brain tissues using Trizol (Ambion). HIV-1 infection was confirmed by detecting the *gag* gene and the *vpr* gene transcription in the Tg26 mice (TG) but not in normal wildtype (WT) mice (**insert**). Transcripts of *TLR4, TNFα, NF-kB*, and *Abcc8/*Sur1 were measured with gene-specific primers. Results are shown in X±SD and analyzed by two-tailed and heteroscedatic t-test: *, *p*<0.05; **, *p*<0.01; ***, p<0.001.

### Correlation of HIV and Vpr with upregulation of Sur1-Trpm4 and host innate proinflammatory signaling in astrocytes of postmortem brain tissues of HIV-infected patients

We next used the same IHC method to examine whether the same set of inflammatory markers are also elevated in astrocytes of HIV-infected human brains. Postmortem brain tissues from HIV-infected and non-infected control subjects in the regions of hippocampus and cerebellum were immunolabeled with the respective antibodies. Consistent with the results from Tg26 mice, Sur1 was significantly elevated (**, *p*<0.01) in HIV-infected astrocytes (**Fig. 3Ab-c**) in comparison with controls (**Fig. 3A-a**), as revealed by the analysis of number of cells double labeled for Sur1 and S100B, a Ca^2+^-binding peptide mainly found in astroglial cells. Furthermore, the Sur1 protein co-localized with Trpm4 (**Fig. 3-A-d**), suggesting upregulation of the Sur1-Trpm4 channel. In agreement with HIV-positive status of these brain tissues, HIV-1 Vpr protein was also highly expressed in some cells (**Fig. 3B**), which correlated with the presence of TLR4, NF-kB (p65) (**Fig. 3C**) and TNF (**Fig. 3D**) in astrocytes. Together, these results showed positive correlations of HIV infection and Vpr with upregulated expression of TLR4, TNFα, NF-kB and Sur1-Trpm4 in astrocytes of both human and mice brain tissues.

**Fig. 3.**
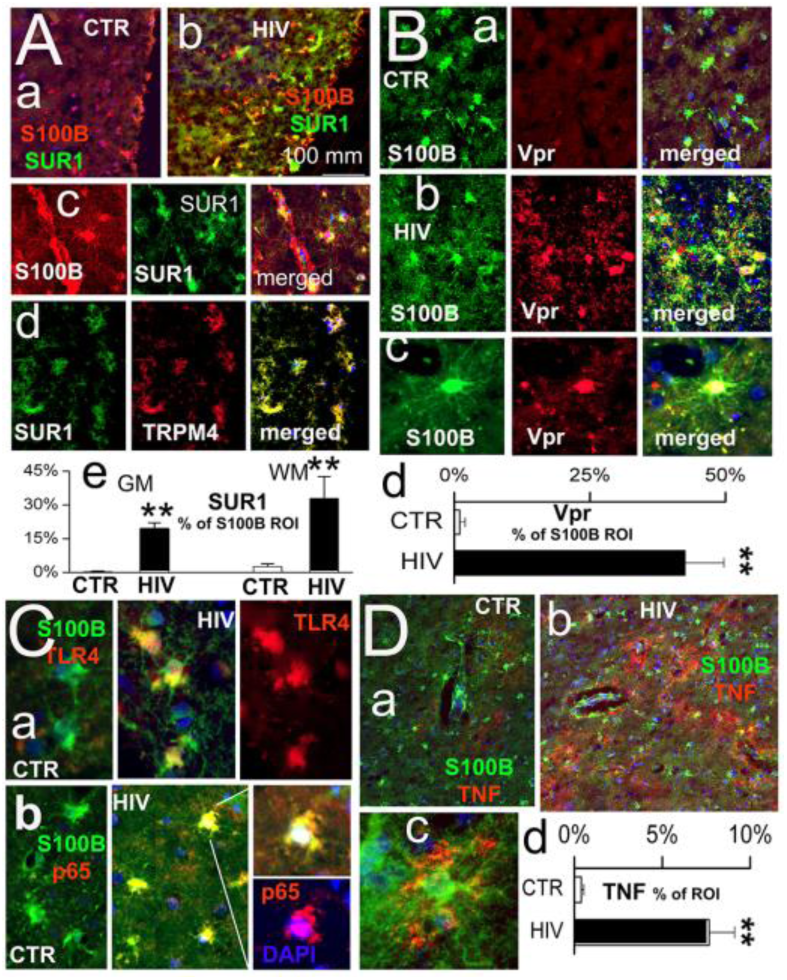
Correlation of HIV and Vpr with host innate immune proinflammatory responses in astrocytes of HIV-infected postmortem human brain tissues. (**A)** Elevation of Sur1, where Trpm4 localizes in HIV-infected astrocytes. (**a**) Control (CTR). (**b**) HIV-infected; (**c**) An enlarged view of (**b**). S100B (red) shows where astrocytes are1. Sur1 (green), and (**d**) co-localization of Sur1 with Trpm4. GM, gray matter; WM, white matter. (**B**) Vpr is highly expressed compared with CTR in astrocytes (**a**) and Vpr (**b-c**). S100B (green), Vpr (red). (**C**) Elevation of TLR4 (**a**) and nuclear NF-kB (p65) (**b**) in astrocytes. DAPI (blue). (**D**) Elevated TNF in astrocytes. Graphs underneath each panel shows % region of interest (ROI). Two-tailed and unpaired t-test: **, p<0.01.

### Expression of HIV-1 *vpr* activates the same set of proinflammatory markers and Sur1 in SNB19 cells

To evaluate the contribution of HIV-1 Vpr protein to the observed upregulation of proinflammatory responsive markers and Sur1-Trpm4 in astroglial cells, we used an adenoviral (Adv) system to deliver Vpr into a human glioblastoma SNB19 cell line with increasing multiplicity of infection (MOI). Mock or Adv-Vpr transduced cells were collected 24 hours (h) post infection (*p*.*i*.). RNA was isolated and subjected to RT-qPCR for mRNA transcriptional analysis. As shown in **Fig. 4**, Vpr increased transcription of *TLR4, TNFα, NF-kB* and *Abcc8/Sur1* genes in a concentration-dependent manner that were statistically significant with the exception of NF-kB due to high experimental variations.

**Fig. 4.**
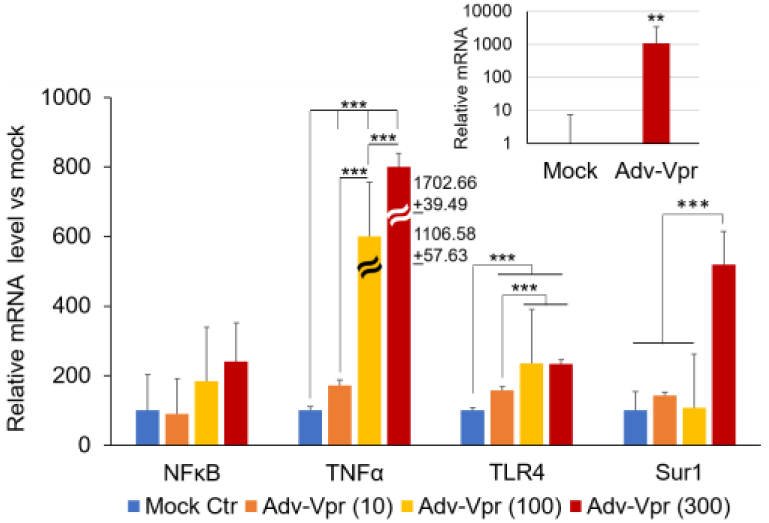
HIV-1 Vpr activates concentration-dependent gene transcription of *TLR4, TNFα, NF-kB*, and *Abcc8* (Sur1) in human glioma SNB19 cells. Adv-Vpr, adenoviral Vpr. Mock or Adv-Vpr transduced cells with MOI of 10, 100 and 300, were collected 24 h post-infection (*p*.*i*.). RT-qPCR was used to measure gene transcription. The insert shows HIV-1 *vpr* gene transcription; whereas no mRNA transcript was detected in mock-treated cells. One-way ANOVA followed by Tukey test was used for statistical comparison analysis. **, *p*<0.01; ***, 12 *p*<0.001.

### Counteraction of Vpr-induced apoptosis by a Sur1 inhibitor glibenclamide in SNB19 cells

We next tested whether Vpr is neurotoxic to a human SNB19 cells. Cells were transduced by Adv-Vpr with increasing MOIs, and cell viability was determined 5 days *p*.*i*. (**Fig. 5)**. While the increase of MOI of Adv control did not cause significant cell death (data not shown), a clear concentration-dependent cell killing was shown as indicated by the trypan blue assay (**Fig. 5A**), which was accompanied by a concentration-dependent decrease of cell viability by the MTT assay (**Fig. 5B**). To further elucidate mechanism of Vpr-induced cell death in SNB19 cells, we further tested the effect of Vpr on apoptosis in SNB19 cells., which was measured by a RealTime Annex V assay (Promega). As shown in **Fig. 5D**, positive control Digitonin indicated proper induction of apoptosis. Mock infection or Adv control showed no or little cell death. In contrast, Adv-Vpr-transduced cells showed dose-dependent increase of apoptosis that was significantly higher than the controls. Thus, these results showed that Vpr induces neurotoxicity *via* apoptosis.

**Fig. 5.**
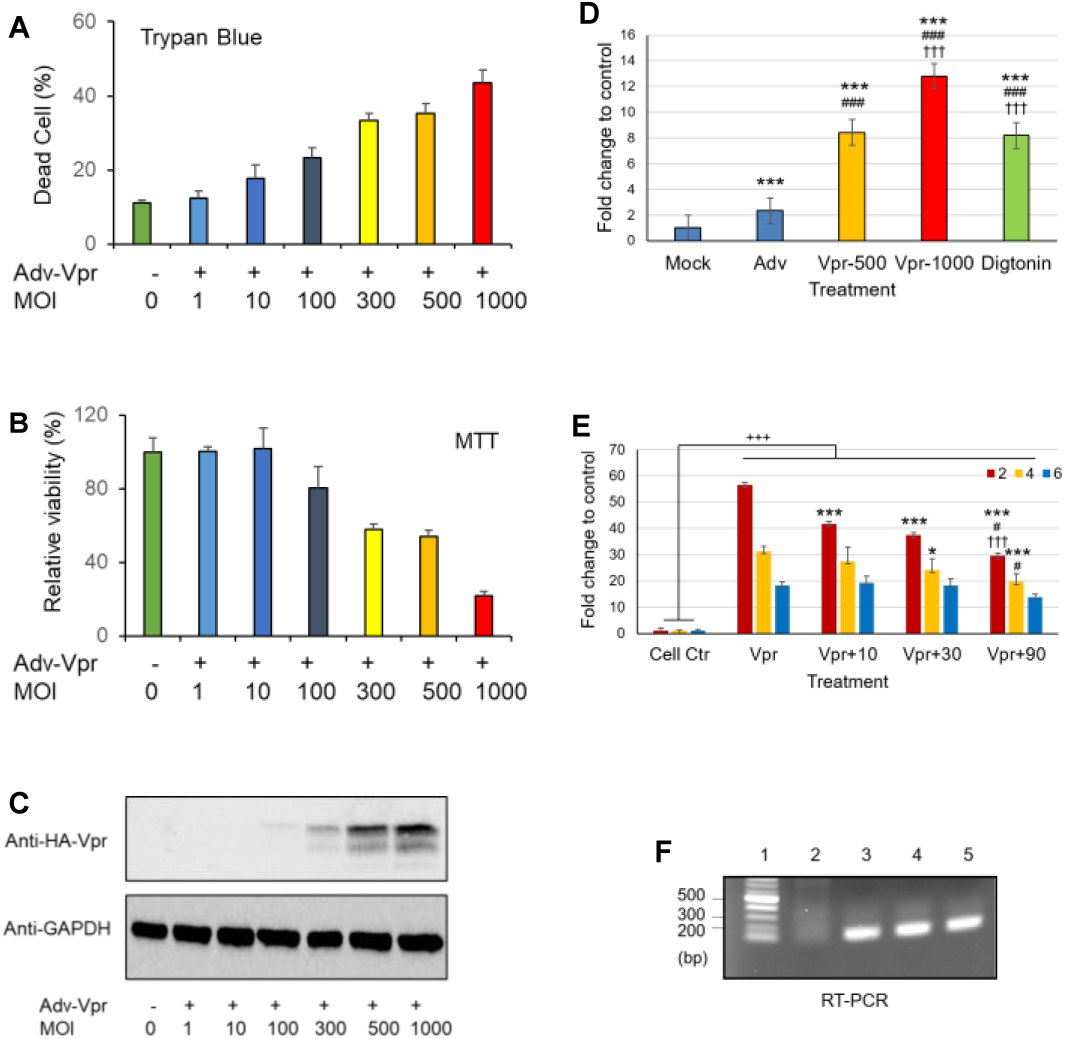
HIV-1 Vpr induces cell death and apoptosis and counteraction by the Sur1 inhibitor glibenclamide in human glial SNB19 cells. HIV-1 Vpr induces cell death and apoptosis in a concentration-dependent manner as measured 5 days *p*.*i*.. by Trypan blue straining (**A**) or by the cell viability MTT assay (**B**). Vpr protein production was confirmed by Western blot analysis (**C**). Note that it was difficult to detect Vpr protein at low MOI. (**D**) Titer-dependent induction of Vpr induces apoptosis. Vpr-induced apoptosis was measured by a RealTime Annex V assay (Promega) 6 h *p*.*i*. Mock, nothing added; Adv, adenovirus control; Digtonin, an apoptosis-inducing agent as a positive control. Data are presented as X±SD. One-way ANOVA followed by Tukey multi-group comparison was used. Treatment: p<0.001; *vs*. mock: ***, p<0.001; *vs*. Adv: ###, p<0.001; *vs*. Vpr-500, †††, p<0.001. (**E**) Concentration-dependent inhibition of Vpr-induced apoptosis by Glibenclamide (10, 30, 90 µM) at 2, 4 and 6 h *p*.*i*. Data are presented as X±SD. Two-way ANOVA followed by Tukey multi-group comparison was used. Treatment: p<0.001; Time: p<0.001; Treatment x Time: p<0.001; *vs*. Vpr: *, p<0.05, ***, p<0.001; *vs*. 30 µM glibenclamide: #, p<0.05; *vs*. 10 µM glibenclamide: †††, p<0.001. (**E**) Vpr gene transcription measured by RT-PCR at 6 h *p*.*i*. Lanes: 1, DNA ladder; 2, Adv control; 3, Adv-Vpr; 4, Adv-Vpr + Glibenclamide; 5, Adv-Vpr positive control. SNB19 cells were infected with Adv-Vpr in a 96-well microtiter plate with MOI as indicated.

Since Vpr increases the expression of Sur1 (**Fig. 4**), we then tested whether a Sur1-specific inhibitor, an FDA-approved drug glibenclamide could suppress the Vpr effect. Vpr-induced apoptosis was measured the same way as shown in **Fig. 5D** after adding increasing concentration of glibenclamide from 10, 30 to 90 µM. A concentration-dependent inhibition of Vpr-induced apoptosis by glibenclamide was observed at 2, 4 and 6 h *p*.*i*. (**Fig. 5E**). RT-PCR analysis indicated that the *vpr* gene was properly expressed (**Fig. 5F**). Together, these data suggest that Vpr induces neurotoxicity *via* apoptosis, which can be alleviated by the Sur1 inhibitor glibenclamide.

## Discussion

Results of this study show the correlation between HIV-1 expression and activation of proinflammatory markers, TLR4, TNFα, and NFκB in astrocytes of HIV-transgenic Tg26 mouse (**Fig. 1-2**) and HIV-infected postmortem human brain tissues (**Fig. 3**), suggesting HIV-1 infection triggers proinflammatory responses of the CNS. In addition, the activation effect of HIV-infection is at least in part due to the presence of HIV-1 Vpr protein because expression of Vpr alone activates the same set of proinflammatory markers as shown in human and mouse brain tissues (**Fig. 4**). Furthermore, the production of Adv-Vpr induces apoptosis in SNB19 cells (**Fig. 5-6**), suggesting Vpr induces neurotoxicity *via* apoptosis. Most interestingly, however, HIV-1 infection or Vpr production is also correlated with the upregulation of the Sur1 and Trpm4 channel (**Fig. 1-4**), and a Sur1 inhibitor glibenclamide inhibits Vpr-induced apoptosis in a concentration-dependent manner in the SNB19 cells (**Fig. 6**). Together, our data suggest that HIV-1 Vpr-induced proinflammatory response and apoptotic cell death are mediated, at least in part, through the Sur1-Trpm4 channel in astrocytes of the CNS.

HAND are a range of neurological disorders of various severity caused by HIV. HIV infects CNS as early as 8 hours after initial infection^22^. It targets primarily human brain in the subcortical areas, and cortex to a less extent, including hippocampus and cerebellum, where it infects glial cells including microglia and astrocytes. Astrocyte is the most abundant type of glial cell. HIV infection of glial cells triggers host proinflammatory immune responses by stimulating reactive astrocytes that release cytokines/chemokines and neurotoxic factors that lead to neurodegeneration, compromised neuronal function and neurocognitive impairments^23^. While cART is effective in eliminating active replicating viruses, it has little or no effect on HIV-1 proviruses that reside in quiescent astrocytes, which therefore become latent reservoirs in patients who otherwise have no detectable or residual HIV-1 under cART^24^. In the absence of active viral replication, chronic HIV infection caused by low levels of viral activities or secretion of neurotoxins of both host and viral origins continue to cause persistent neuroinflammation with neurotoxicity leading to chronic HAND^5^. Consequently, the severity of some HAND does not always directly correlated with the level of HIV replication or viral load, but rather with lasting glial activation^7^, suggesting other HIV-associated factors but not the whole virus *per se* contribute to those HAND. Results of this study provides strong support that HIV-1 Vpr is one of those HIV-associated proteins contributing to HAND.

Vpr could exert its effects on the CNS both as an intracellular and an extracellular protein. As an intracellular protein, it involves in in establishing and sustaining CNS infection, as its function is required for HIV-1 replication in non-dividing monocyte-derived macrophages (MDM), a key lineage of cells involved in HIV-1 neuroinvasion^25^. Vpr can also be found as an extracellular and free circulating protein in blood, cerebrospinal fluid (CSF) and CNS-associated cells^8,13,26-31^, where Vpr reactivates HIV replication from latency^11,26,27,30^ (for detailed reviews, see^8^). Vpr triggers proinflammatory reactions^8,13,29^, and promotes astroglial secretion of proinflammatory cytokines IL-6^32^. Vpr is a also neurotoxin that induces apoptosis in astrocytes^29^, neurons^10^, and in CNS^33^. Consequently, Vpr causes neurodegeneration^10^, synaptic loss and neurocognitive impairment^34^ that link to various HAND^12,13^. In consistent with the literature, our results support a specific role of Vpr-induced neuroinflammation and neurotoxicity in HAND.

One of the most interesting findings of this study is that we discovered a connection between Vpr and the Sur1-Trpm4 channel. Sur1 (sulfonylurea receptor 1) and Trpm4 (transient receptor potential melastatin 4) are two subunits of the Sur1-Trpm4 cation channel. The regulatory subunit Sur1, encoded by the *Abcc8* gene, is a member of the ATP-binding cassette protein superfamily; and the pore-forming subunit Trpm4 is encoded by the *Trpm4* gene. Association of Sur1 and Trpm4 forms heterodimers and a functional Sur1-Trpm4 channel that antagonizes calcium influx mediated by receptor operated calcium entry (ROCE) cation channels^35^. This ion channel is not expressed constitutively but is transcriptionally upregulated in microglia and astrocytes^36^ in responsive to various neuroinflammatory brain conditions including traumatic brain injury (TBI)^37^, subarachnoid hemorrhage (SAH)^38-40^, and neuroinflammatory conditions such as multiple sclerosis (MS)/experimental autoimmune encephalomyelitis (EAE)^21,41^. Thus, the Sur1-Trpm4 channel is a key neuro-regulator involved in various neurological disorders including brain injuries and neurodegeneration^5,20^. Furthermore, the link between Vpr and the Sur1-Trpm4 channel is functionally relevant because Vpr activates NFAT through Ca2+ influx in neuronal SH-SY5Y cells^11,13^. Therefore, it is possible that Sur1-Trpm4 channel-induced neuropathological effects are responsible for the Vpr effects or are part of the collective induction of Vpr-induced HAND. If this is true, this finding might be clinically significant because the Sur1-Trpm4 channel has shown to be a key drug target of various neuroinflammatory conditions^21,37-41^, and pharmacologic inhibition of this channel by the repurposed and FDA-approved drug glibenclamide significantly improved clinical outcomes of those neurologic disorders^21,42-45^. Indeed, the Sur1 inhibitor glibenclamide suppresses Vpr-induced apoptosis in a concentration-dependent manner (**Fig. 6**). Altogether, we may have uncovered a novel HAND-underlying mechanism involving Vpr-induced activation of the Sur1-Trpm4 channel. Once substantiated, it would allow us to test the possibility of treating HAND-afflicted individuals by using the FDA-approved Sur1 inhibitory drug glibenclamide, which has already been shown to be an effective drug for treatment of various neurological disorders^21,42-45^.

## Acknowledgement

The authors would like to thank Dr. Thomas Blanchard of University of Maryland Brain and Tissue Bank for the postmortem human brain tissues, Dr. Joseph Bryant of the Institute of Human Virology, University of Maryland School of Medicine for the Tg26 mouse brain tissues, and Dr. Hengli Tang of Florida State University for the SNB19 cell line. This research was support in part by an intramural support from the University of Maryland Medical Center and a VA merit-based award (I01 BX004652 to RYZ) and). V.G is supported by grants from NINDS (R01NS061934 and R01NS107262), and J.M.S. is supported by grants from the NHLBI (R01HL082517) and NINDS (R01NS060801, R01NS102589, and R01NS105633).

